# Kinematics associated with treadmill walking in Rett Syndrome

**DOI:** 10.1101/568360

**Authors:** Charles S. Layne, David R. Young, Beom-Chan Lee, Daniel G. Glaze, Aloysia Schwabe, Bernhard Suter

**Affiliations:** Health and Human Performance, University of Houston, Houston, Texas, United States of America; Center for Neuromotor and Biomechanics Research, University of Houston, Houston, Texas, United States of America; Center for Neuro-Engineering and Cognitive Science, University of Houston, Houston, Texas, United States of America; Texas Blue Bird Circle Rett Center, Texas Children’s Hospital, Houston, Texas, United States of America; Baylor College of Medicine, Houston, Texas, United States of America

## Abstract

Individuals with Rett syndrome suffer from severely impaired cognitive and motor performance. Current movement-related therapeutic programs often include traditional physical therapy activities and assisted treadmill walking routines for those patients who are ambulatory. However, there are no quantitative reports of kinematic gait parameters obtained during treadmill walking. Here we report the results of an investigation of 17 females diagnosed with typical Rett who walked on a treadmill as speed gradually increased. The objective included characterizing lower limb kinematics, including knee and hip joint range of motions, velocities, limb asymmetries, and the variance associated with these measures. Joint kinematics were obtained using a 12 camera motion capture system and associated processing and analysis software. Stride times progressively decreased as treadmill speeds increased although the range of speeds our participant could walk was quite slow: range 0.2 m/s – 0.5 m/s. There were significant main effects of speed on sagittal knee and hip range of motions and hip velocity. There were large joint asymmetries and variance values relative to both healthy walkers and others patient populations although variance values decreased as walking speed increased. There were significant correlations between joint range of motions and stride times and joint velocities and stride times. The results indicate that Rett patients can adapt their kinematic gait patterns in response to increasing treadmill speed but their ability to do so lies within a narrow range of speeds. We suggest that treadmill training for ambulatory individuals with Rett may further promote improved walking kinematics as well as overall health benefits.

## Introduction

Mutations in the gene coding for methyl-CpG-binding protein 2 (*MECP2*) result in the neurodevelopmental disorder Rett Syndrome (RTT). Although a relatively rare condition, worldwide RTT affects approximately 1 in 10,000 live born females [1]. Seemingly normal development occurs up to ages 6-18 months at which time a period of regression with loss of verbal skills and social interactions, as well as both fine and gross motor skills sets in. Stereotypical hand movement, breathing difficulties, apraxia, ataxia, muscle hypertonia, limb rigidity and bruxism are some of the disabling symptoms commonly observed. A period of stabilization ensues, but bipedal postural control and walking are severely compromised and walking ability often declines further at later ages, such that ultimately less than half remain able to walk.

Loss of ambulatory skills results in a number of additional physical problems such as muscle atrophy, limb contractures, decreased cardio-respiratory fitness and low overall physical fitness. Suggested physical therapy for patients with RTT has included physical exercise designed to increase physical fitness and to maintain walking ability. These therapies have ranged from traditional physical exercises and stretching [2] guided physical activities in a multi-sensory room [3], and hydrotherapy [4]. Other authors have suggested that a program incorporating walking may have a range of benefits for individuals with RTT including improved physical fitness as well as positively influencing their quality of life and wellbeing [5].

There are several reports of those with RTT exploring the possibility of incorporating treadmill walking into their therapeutic regimen. Lotan et al. [6], explored the use of a treadmill-walking program to promote both walking skills and physical fitness and reported high correlations between improved walking performance and physical fitness. While promising, this study was conducted with only four girls with RTT. Three girls with RTT served as subjects for an exploratory investigation using robot-assisted walking [7] with preliminary results indicating the girls tolerated the robotic system suggesting that this might be a clinical tool that merits further investigation with RTT patients. These preliminary studies, while promising, leave open the applicability of these findings to a broader spectrum of ambulatory girls with RTT.

Motorized treadmills have been used extensively for gait training with populations suffering from conditions such cerebral palsy, stroke, and Parkinson’s disease. The results of these studies indicate improvement in gait patterns after treadmill gait training. Gait training with motorized treadmills is standard practice with a variety of populations with gait disorders, including those with Parkinson’s disease, cerebral palsy and stroke. Multiple investigations have documented improvements in overground gait parameters with the aforementioned populations resulting from treadmill gait training [8-13]. Individuals with RTT have neurological factors that are generally different than of those conditions listed immediately above, however there is no a priori reason to suggest that treadmill training would not provide benefits to those with RTT. Currently, treadmill walking is often a component part of therapy programs for those with RTT however its efficacy is unknown.

Prior to exploring the efficacy of a treadmill-walking program for both improved overall physical fitness and functional walking characteristics, it is important to identify some typical kinematic features associated with treadmill walking of patients with RTT. Such information is necessary to evaluate any potential improvements stemming from a treadmill-walking program. Although Downs and her colleagues have completed extensive efforts to develop reliable and valid measures of RTT walking that can be used as clinical measures [14] currently there are no reports of lower limb quantitative kinematics obtained from individuals with RTT during treadmill walking.

An important characteristic of effective walking is the ability to adapt to different speeds. Therefore, we were interested in determining if RTT patients could modify their gait to keep pace with increases in the speed of the treadmill. The basic rhythmical, alternating limb pattern driving locomotion has long been proposed to be the product of a network of spinal neurons that require the mediation of higher order structures for the complete expression of goal-directed walking. This network is commonly referred to as a central pattern generator i.e. CPG [15, 16, 17]. Tonic innervation of the spinal locomotion circuits is regulated by noradrenalin and serotonin neurons. Without this innervation, which [18] suggests is impaired in those with RTT, due to the hypofunctioning of the aminergic neurons in the brainstem, proper functioning of the spinal circuit is impaired. However, input into the circuit from lower limb muscle spindles and foot contact information [19] can assist in activating the circuit to generate the basic locomotor pattern [20]. The well-documented toe walking exhibited by RTT patients is proposed to be an adapted behavior that generates increased spindle input to the spinal circuitry and therefore activates the circuit [18]. This suggests that the basic spinal locomotion circuity remains generally intact and can be activated with increased sensory input.

Successful adaptation to increasing treadmill speed for those with RTT would indicate that the neural mechanisms available to integrate the peripheral sensory information associated with increased limb speed in a manner that lower limb kinematic parameters could be successfully adapted to walk at a faster speed. Previous work by our group explored details of the temporal features of gait of individuals with RTT during both overground and treadmill walking [21]. It was reported there were increases in stance time but decreases in swing and double support time when comparing treadmill to overground gait. Additionally, treadmill walking resulted in decreased variance in the temporal gait parameters, indicating treadmill walking resulted in a more regularized gait. The current work provides the first description of quantitative kinematic data obtained during treadmill walking as the speed of the treadmill progressively increased.

## Methods

### Study Participants

Seventeen females diagnosed with typical RTT based upon the Neul et al. [22] criteria and carrying pathogenic MCEP2 mutation served as subjects in this study. They ranged in age from 4 to 20 with a mean age of 10.8, standard deviation ± 5.3 and were receiving treatment at the Blue Bird Circle Rett Center at Baylor College of Medicine in Houston, TX. All subjects were able to independently walk without orthotics and none were taking medication that would be expected to impact their motor control function including benzodiazepines (often used for muscle tone control). All procedures were approved by the Institutional Review Boards of the Baylor College of Medicine (H-35835) and the University of Houston (00000855). The parents provided written informed consent for their daughters.

### Data Collection

The task involved the subjects walking on a duel-belt motorized treadmill (Bertec^®^) that contained force plated embedded under each belt. The subjects were secured in an overhead harness that eliminated any potential falls but did not provide postural support during walking. Walking was initiated at 0.1 m/s and was increased by 0.1 m/s every 20 seconds until either the parents indicated that was the maximum speed the subject could obtain or the subject began to exhibit signs of discomfort such as vocalizations, hand or facial gestures. Depending upon the subject’s gait pattern and treadmill speed, the 20 seconds of data collection resulted in 10-14 strides for each treadmill speed.

Kinematic data were collected at 100 Hz using a Vicon^®^ 12-camera motion capture system in combination with the plug-in gait data processing software. Reflective markers were applied bilaterally on the heel, toe, ankle, knee, shank and hips prior to data collection. Ground reaction forces from the treadmill force plates were sampled at 1000 Hz and synchronized with the kinematic data. Kinematic and force data were used in combination to identify heel strike and toe off. Additional details regarding the data collection procedures can be obtained in Layne et al. [23].

### Data Processing and Analysis

A preliminary assessment of the data revealed that all 17 subjects were able to walk between the speeds of 0.2 and 0.5 m/s therefore the decision to analyze the kinematics associated with the speeds of 0.2, 0.3, 0.4 and 0.5 m/s was made. A custom MATLAB (MathWorks^®^) was used to filter the kinematic data with a Butterworth low-pass filter with a 6 Hz cut-off frequency.

Bilateral heel strikes were detected and the data between consecutive ipsilateral heel strikes were saved as individual strides for both the right and left legs. Heel strikes were identified at the minimum position of the heel marker during each gait cycle. The toe marker minimum was used in the event the subject was toe walking on particular strides. The kinematic data were then time normalized such that each stride was represented by 100 samples. The time normalized waveforms were then amplitude normalized such that the angular value heel strike was zero degrees. For each normalized stride, sagittal plane knee and hip angles were obtained for each treadmill speed, for each subject. Maximum and minimum angular values were obtained and used to calculate the range of motion (ROM) for each stride. After the individual joint angles were obtained, the velocity curves for each angle were calculated. Peak angular velocity for each stride and each subject were also identified.

After the above processing was completed, the limb with the greater ROM, for each joint, was identified. The data was then reorganized into the side (i.e. left or right) with the strides of the greater ROMs grouped together and those stride with lesser ROMs grouped together. Symmetry indexes (SI) between greater and lesser joint angles were computed using the following formula [24]. A SI of 0 reflects perfect symmetry between the two limbs.

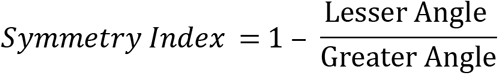

After it was determined that there were no significant differences between the joint ROM and associated peak velocities, the data from the two limbs was collapsed for further processing and analysis. The data from each variable were then averaged for each subject, at each gait speed, and group means calculated. It was found that many of variables were not normally distributed based upon the results of the Shapiro-Wilk test of normality. Therefore, Friedman tests were used to determine if significant differences existed between the ROMs for each joint across the four treadmill speeds. Follow up Wilcoxon tests were utilized as appropriate with a Bonferroni correction being applied. An alpha level of p < 0.05 was adopted for significance. Pearson’s correlation coefficients between a joint’s ROM and its velocity and between stride times and ROMs were calculated. The above procedures were also applied to the peak velocity values to determine if limb velocity changes in response to increasing treadmill speed. To determine the relationship between the various variables associated with the gait of individuals with RTT, Pearson correlations coefficients between a joint’s ROM and its velocity, between stride times and ROMs and between stride times and joint velocities were calculated. Correlations were also developed between subject age and joint ROMs and velocities. Finally, correlations were developed between stride times and subject age. To assess if the variance of the dependent measures was influenced by treadmill speed, the *F* test for equality of variance was employed.

Occasionally our subjects’ feet would cross the midline and land with one foot in front of the other. Therefore, we were interested in determining the degree of knee joint motion in the horizontal plane. We applied the same processing techniques for the knee motion in the horizontal plane as those used for sagittal joint angles. Additionally, Downs et al. [5] reported minimal vertical motion of the hip during overground walking in her subjects with RTT assessed with the Actigraph GTX3 tri-axial accelerometer device. To determine if this reported lack of vertical hip motion is a common feature of RTT gait, we analyzed the motion of the pelvis in the coronal plane. Based on the literature, we identified the range (plus two standard deviations) of transverse knee motion for healthy individuals and determined which of our subjects exceeded that range. Similarly, we identified the range (minus two standard deviations) of the vertical motion of the hip and determined if any of our subjects failed to reach the degree of motion demonstrated by healthy walkers. Descriptive statistics of the number of subjects who either exceeded the healthy range of transverse knee motion or failed to display the healthy amount hip vertical motion are reported.

## Results

The primary purpose of this investigation was to determine if individuals with RTT were able to adapt their lower limb kinematics and associated stride times as treadmill speed progressively increased. Secondary considerations included exploring the prevalence of excessive knee joint motion in the horizontal plane and pelvis motion in the frontal plane.

Table 1 displays that as treadmill speed increased from 0.2 to 0.5 m/s, our subjects were able to decrease their stride times so they could continue walking. The Friedman test revealed a significant effect for speed (χ^2^ = 86.698, p < 0.000). However, only three of the 17 subjects tested were able to continue walking up to the speed of 0.6 m/s. Thus, although our subjects were able to adapt to the increasing treadmill speeds, that ability was limited to a narrow range of speeds.

**Table 1-.** Median stride times by speed and statistical comparisons

| Table 1- Median stride times by speed and statistical comparisons |  |  |  |  |
| --- | --- | --- | --- | --- |
| Speed | Median (s) | Comparison | Z value | P values |
| 0.2 | 1.43 |  |  |  |
| 0.3 | 1.27 | 0.2 vs 0.3 | -4.525 | 0.000 |
| 0.4 | 1.21 | 0.3 vs 0.4 | -4.505 | 0.000 |
| 0.5 | 1.17 | 0.4 vs 0.5 | -4.368 | 0.000 |

### Joint ROMs

The Friedman test revealed a significant main effect of speed on sagittal knee ROM (χ^2^ = 11.047, p < 0.011). Follow up Wilcoxon tests indicated that the ROM between speeds 0.2 and 0.3 (0.2 median = 9.425, 0.3 median = 10.03, Z = −2.812, p < 0.005) and 0.2 and 0.4 significantly different (0.2 median = 9.425, 0.4 median = 9.99, Z = −2.445, p < 0.014). No other comparisons reached significance (Figure 1). Comparisons between the sagittal hip ROM and treadmill speed revealed a significant main effect of speed ((χ^2^ = 14.012, p < 0.003). Significant ROM differences existed between the ROM for speeds 0.2 and 0.3 (0.2 median = 10.155. 0.3 median = 11.385, Z = −3108, p < 0.003). There were no other significant differences for the hip ROM comparisons.

**Figure 1.**
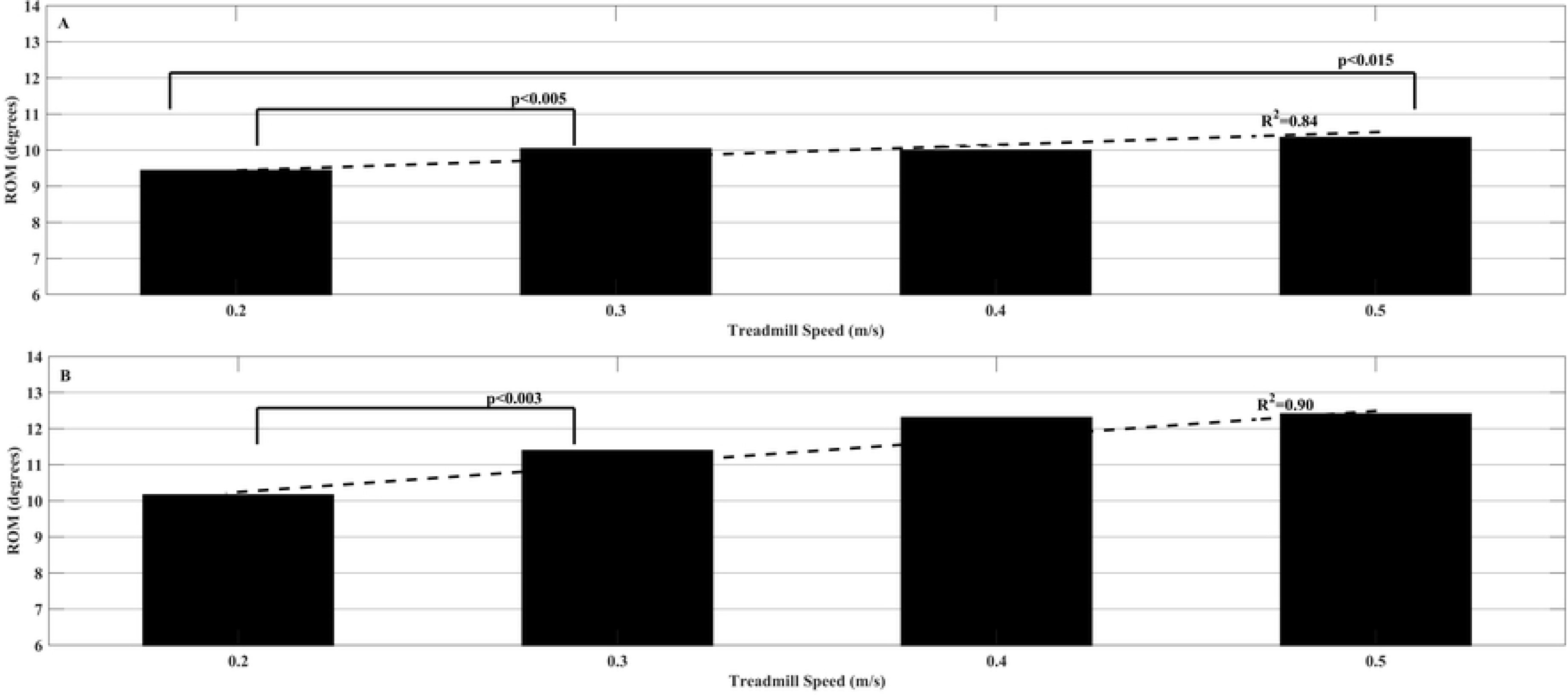
Median Knee (A) and Hip (B) ROM across treadmill speeds

### Joint Peak Velocities

For peak sagittal knee velocities in degrees per second, the Friedman test approached significance (χ^2^ = 7.238, p <0.065). The Friedman test for peak sagittal hip velocities revealed a significant main effect of treadmill speed (χ^2^ = 14.633, p < 0.002). There were significant increases between treadmill speeds 0.2 and 0.3 (0.2 median = 35.0, 0.3 median = 39.1, Z = −2.711, p < 0.007) between speeds 0.2 and 0.4 (0.2 median = 35.0, 0.4 median = 45.3, Z = −2.711, p < 0.007). Interestingly there was a significant decrease between speeds 0.4 and 0.5 (0.4 median = 45.0, 0.5 median = 39.5, Z = −2.744, p < 0.006). The Friedman test for the knee velocity in the horizontal plane revealed no significant differences across the four speeds (χ^2^ = 4.575, p < 0.206). Figure 2 displays the median angular velocities across the treadmill speeds and the associated R^2^ values.

**Figure 2.**
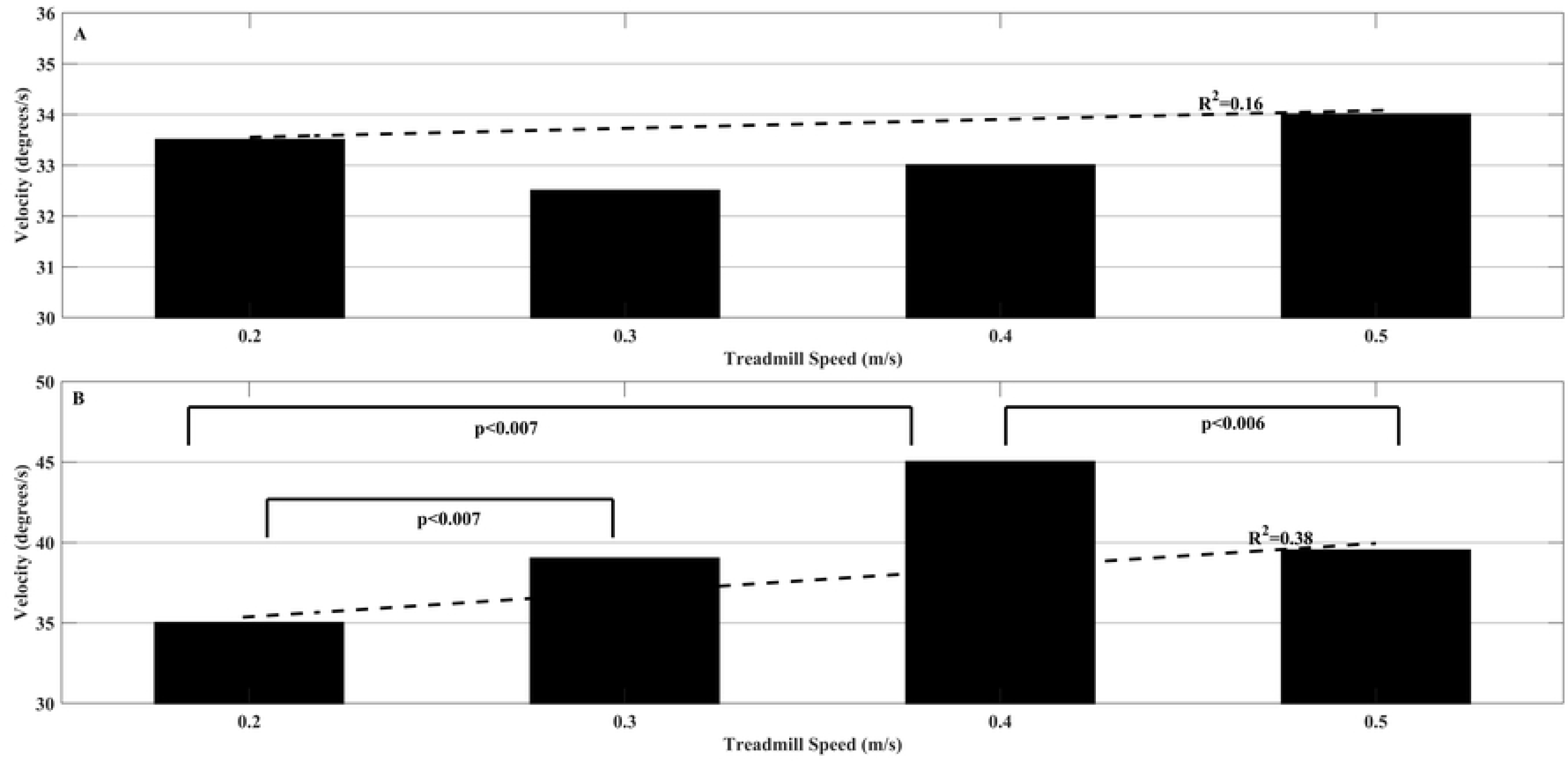
Median knee (A) and hip (B) joint angular velocity across speeds.

There were no significant changes in the SIs for the knee and hip in the sagittal plane. Figure 3 does reflect that our subject’s gait was asymmetrical with all SI values being significantly greater than 0 (i.e. perfect symmetry).

**Figure 3.**
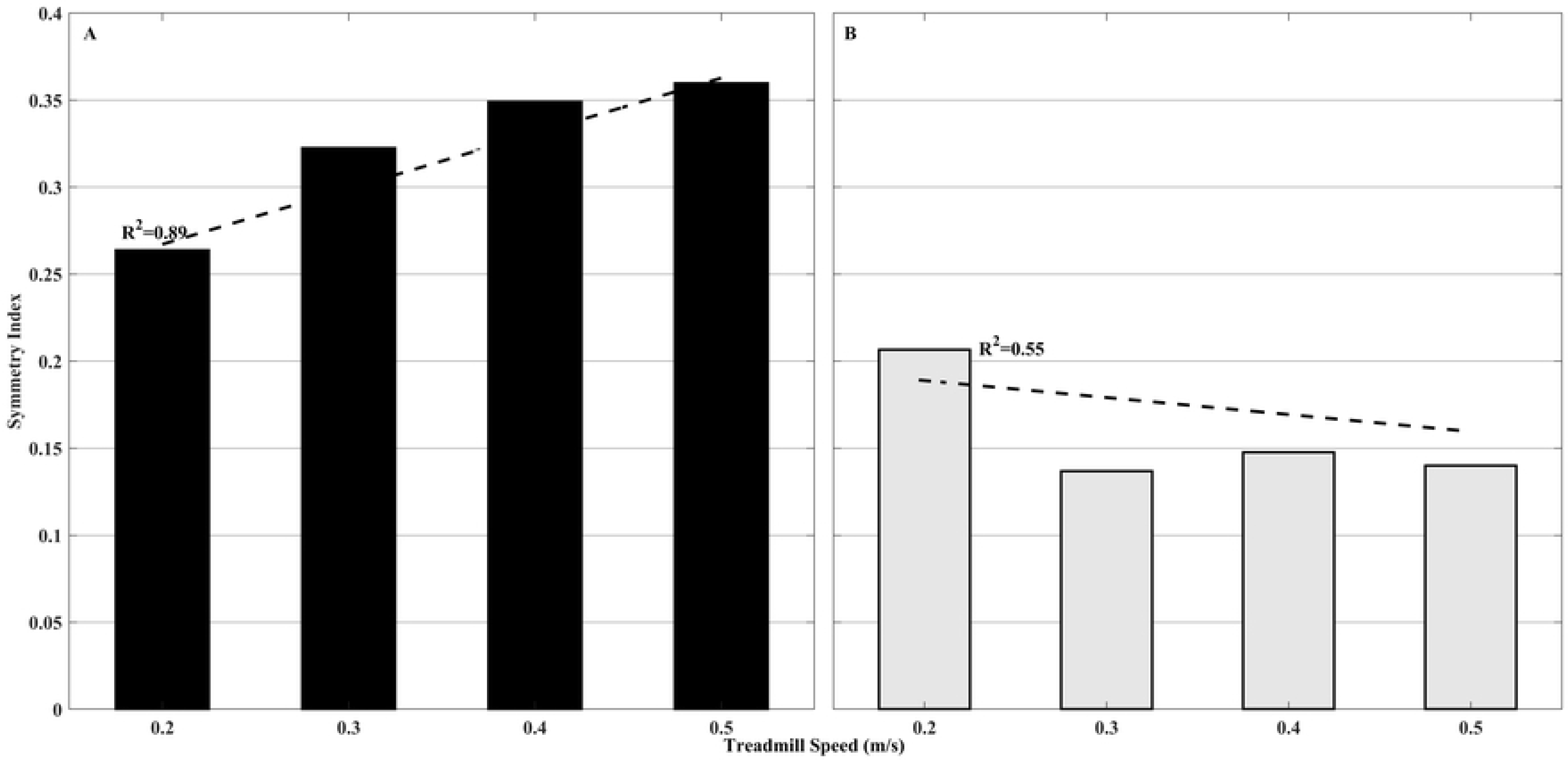
Symmetry index values for the sagittal plane motion knee (solid fill) and hip joints across treadmill speeds.

The Pearson correlation value between subject age and stride time was 0.46 (R^2^ = 0.21) which is significant at the p < 0.01 level. Figure 4 illustrates the high correlations between the joints’ ROM and their associated velocities across the treadmill speed increases.

**Figure 4.**
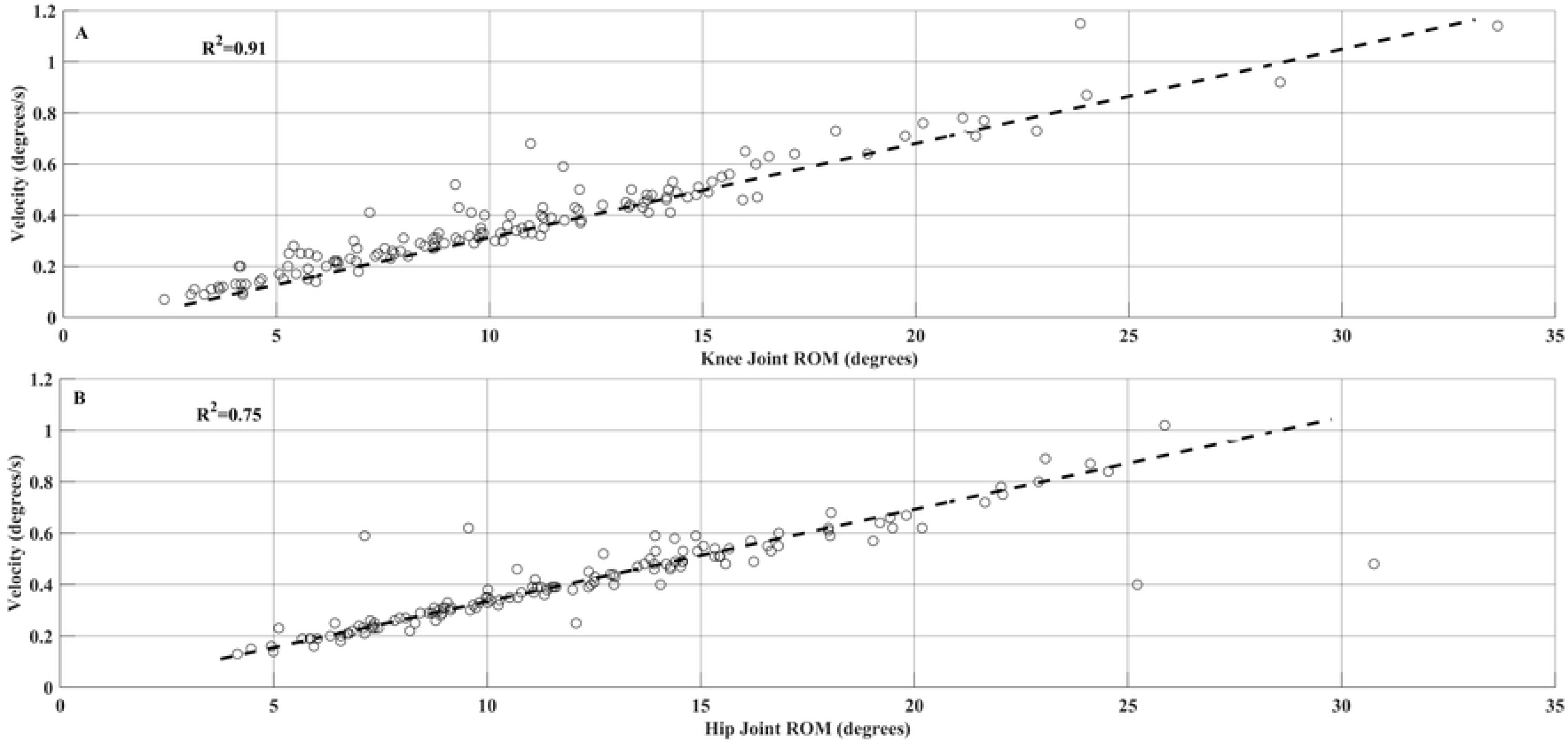
Relationships between ROM and associated angular velocities across treadmill speeds.

Table 2 displays the Pearson R coefficients of the comparison between kinematic variables, stride times and subject age.

**Table 2 –.** Correlation between stride times, kinematics and age

| Table 2 – Correlation between stride times, kinematics and age |  |  |
| --- | --- | --- |
| Pearson Correlation Coefficients (R) | S Knee | S Hip |
| Joint ROM & Stride Times | 0.38* | 0.59* |
| Joint Angular Velocities & Stride Times | 0.45* | 0.60* |
| Joint ROM & Age | 0.15 | 0.37* |
| Joint Angular Velocity and Age | 0.12 | 0.36* |

\*Significant at $p < 0.01$

The F tests to assess potential differences in the variance associated with the joint ROM across speeds indicated that although the variance values were very high, there were no differences resulting from changes in treadmill speed. The same was true for the F tests comparing knee velocities across speeds. However, significant differences in variances were found for hip velocities between speeds 0.2 vs 0.4 (F = 2.444, p < 0.006), 0.2 vs 0.5 (F = 3.292, p <0.000), and 0.3 vs 0.5 (F = 2.169, p < 0.015). In all cases of significant F tests, the slower speed was always associated with the greater variances relative to the faster speed (Table 3).

**Table 3.** Variance values for sagittal plane ROM, and joint velocities

| Table 3. Variance values for sagittal plane ROM, and joint velocities |  |  |  |  |
| --- | --- | --- | --- | --- |
| Speed | Knee ROM | Hip ROM | Knee Velocity | Hip Velocity |
| 0.2 | 25.9 | 32.0 | 0.048 | 0.071 |
| 0.3 | 35.8 | 30.3 | 0.046 | 0.047 |
| 0.4 | 34.8 | 23.1 | 0.043 | 0.029 |
| 0.5 | 21.7 | 19.9 | 0.027 | 0.022 |

Using a ROM of 10° to indicate excessive motion in the transverse plane based on values obtained with healthy individuals [25,26,27], there were only 13 instances, of a possible 136 that exceeded that threshold across all speeds and the two legs. There was no systematic effect of either age or speed on knee transverse plane motion. To determine if our subjects displayed healthy pelvic motion in the coronal plane we, used a threshold of 0.7° as a minimum value to indicate if there was adequate peak motion in this plane [28,29]. Of the 136 measures, only six values fell below the minimal threshold value and these values were confined to just two subjects. These data confirm, with very few exceptions, our subjects with RTT displayed a range of hip motion in the coronal plane associated with healthy gait. The median transverse plane knee ROMs and median peak degrees of the hip in the coronal plane across speeds are displayed in Figure 5.

**Figure 5.**
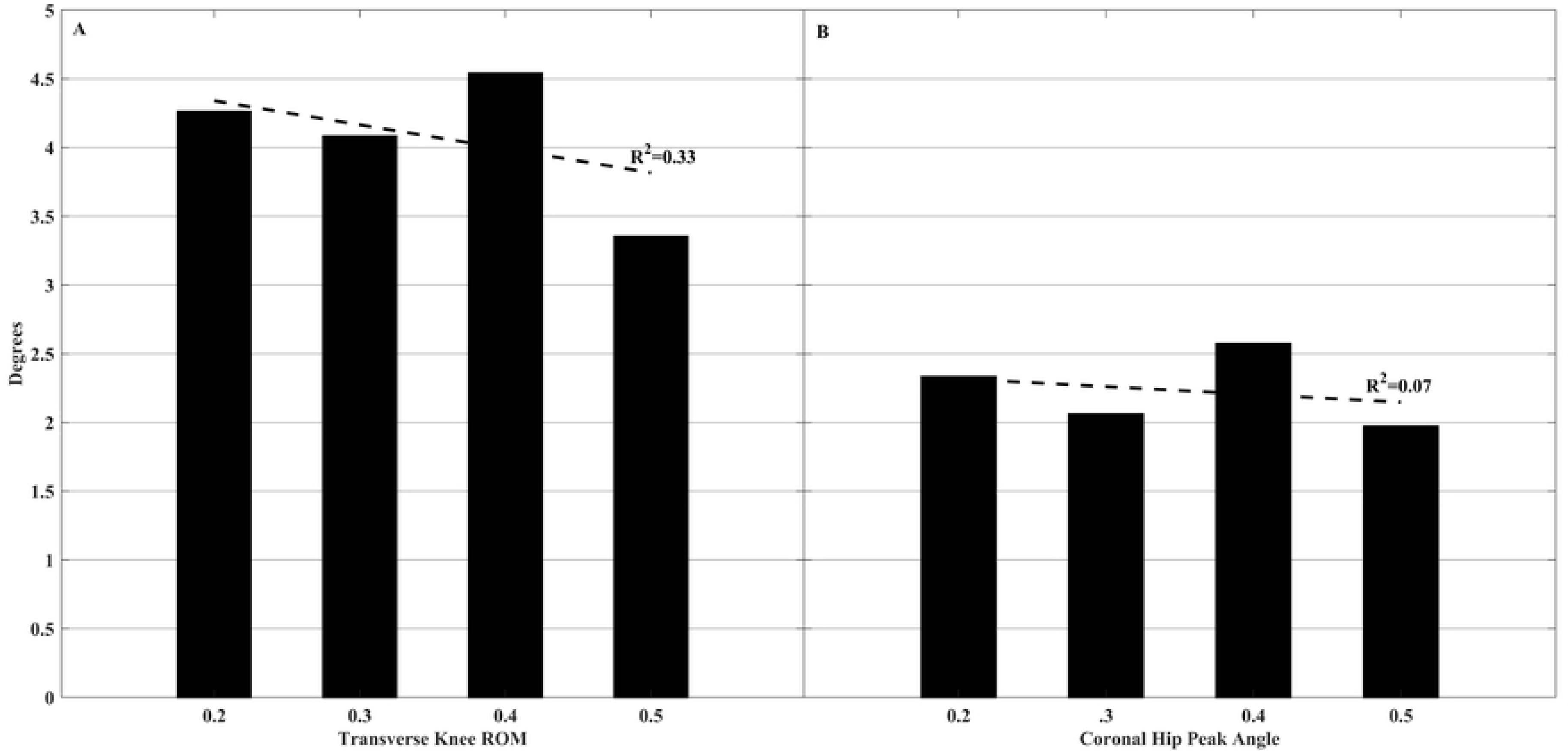
Median ROMs for knee transverse plane motion and median peak degrees for hip coronal plane motion across treadmill speeds.

## Discussion

In this report, we provide the first laboratory-based information regarding kinematic gait data collected from females with RTT. Characterizing the kinematic parameters associated with walking of patients with RTT is important to determine if pharmacological or therapeutic approaches are successful. Additionally, we were interested in determining if those with RTT were able to successfully adapt their gait to increasing treadmill speeds. If so, this would suggest that despite abnormal kinematic parameters, neurological mechanisms remain intact to respond to the sensory feedback associated with increased treadmill speed and adapt their kinematic parameters accordingly.

As reported in Table 1, the subjects were able to decrease their stride times as treadmill speed increased as has been demonstrated in a sample of healthy subjects [30]. However, these decreases occurred within a relatively narrow range of treadmill speeds (0.2-0.5 m/s). To place both the treadmill speed and the stride times in perspective, in a large study of typically developing children ranging in age from 5 to 12, Lythgo et al. [31] found that when a sample of children who averaged 5.7 years of age were asked to walk slow, they averaged 0.97 m/s with average stride times of 0.99 seconds. Our average stride times ranged from 1.45 seconds at speed 0.2 m/s to 1.10 seconds at speeds 0.5 m/s. The average 10.5 year old (similar to the average age in this investigation) in the Lythgo study averaged 1.04 m/s with average stride times of 1.15 seconds when asked to adopt a slow gait. To provide additional perspective, 9.5 year old children diagnosed with spastic diplegic cerebral palsy (CP) walked at a self-selected speed 0.86 m/s on average during overground walking [32]. This value is 72% greater than the maximum speed our patients walked on the treadmill. An additional study reported that 10 year old children with bilateral CP walked at 0.83 m/s on average while those with unilateral CP walked on average at 1.01 m/s [33]. Although not particularly surprising, these comparisons between children with CP and our subjects of similar age emphasize that girls with RTT walk significantly slower than those with CP.

Despite the minimal range of slow walking speeds, our subjects did decrease their stride times such that they were able to maintain pace with the increasing treadmill speeds. This finding strongly suggests that our subjects were able to both adequately detect the sensory information indicating the treadmill speed was increasing and integrate that information to increase their lower limb velocities that resulted in significantly decreased stride times. This is consistent with Aoi et al’s [19] assertion that foot contact information and muscle spindle input can activate the CPG and adjust the locomotor pattern to meet the lower limb movement demands associated with increasing treadmill speed. Consistent with the decrease in stride times are the significant increases of knee and hip ROMs and angular velocity associated with increases treadmill speed as has been reported for a large range of healthy individuals [34,35,36]. These significant main effects and the highly significant correlations between knee and hip ROM and their associated angular velocities (see Figure 5) also reflect our subjects’ ability to modify their lower limb kinematic motion to adapt to the increasing treadmill speeds. Our data thereby suggest that our females with RTT do have intact spinal locomotion circuity that can be regulated by the sensory input generated by walking within a narrow range of walking speeds. We speculate that our subjects are unable to increase their walking speed beyond 0.5 m/s is primarily related to their failure to maintain their attention on the walking task as well as their inability to preserve postural stability despite the safety that the harness provided.

Although the spinal CPG may be able to produce the fundamental alternating lower limb motion necessary to walk, the associated kinematics display a large amount of variance and the relationship between the two limbs is asymmetric. These features contribute to our subject’s lack of postural stability, which therefore prevents them from being able to increase their walking speed. As observed in Figures 3 and 4, our subjects had large symmetry indices and it is worth noting that there was a significant linear trend for knee flexion asymmetry to increase as treadmill speed increased (R^2^ = 0.90). For comparative purposes, a gait study of patients with peroneal nerve palsy displayed median knee joint angular asymmetry of 20% from perfect symmetry [37] while a group of healthy subjects displayed a 3.7% deviation from perfect symmetry [38]. In contrast, our knee joint asymmetries ranged from 26% at 0.2 m/s to 36% at 0.5 m/s. reflecting a high degree of asymmetry.

Another notable feature of the kinematics exhibited by our subjects is the very small range of knee joint motion despite some minimal but statistically significant speed-related increases. Consistent with our results, previous investigations have also reported minimal changes in knee motion associated with small increases in walking speed [39]. The median values ranged from 9.4 at 0.2 m/s to 10.3 at 0.5 m/s. This minimal knee ROM can be characterized as ‘stiff-knee gait’ (SKG) and contributes to the slow speeds at which our subjects were able to walk. °. Healthy individuals when asked to walk at 0.3 m/s on a treadmill, a speed that our subjects walked, had an average knee ROM of 46.1 and a hip ROM of 29.6 [39]. Carriero et al. [32] published data from a sample of children with spastic diplegia CP aged 9.5 years and reported a mean range of 41.3° while an aged match sample of typically developing children displayed a ROM of 65.4°. Individuals post-stroke also exhibit significantly reduced knee ROM during gait [40,41]. For example, the post-stroke subject’s in Chen et al.’s [40] investigation displayed peak knee flexion of 37.8° with their paretic limb while healthy controls had average peak knee flexion values of 61.9. Thus, even patient populations that have been characterized as displaying SKG had significantly greater knee motion that ambulatory females with RTT. Concerning hip ROM in the sagittal plane, Carriero’s et al. study [32] reported a range of 47.1 for children with CP and 49.9 for typically developing children. Again, these values are significantly greater than observed in the current study.

Our sample of females with RTT have an extremely limited lower limb ROM as well as a limited range of walking speeds. Both post-stroke individuals and those with CP who exhibit stiff-knee gait also display compensatory kinematic strategies, primarily hip hiking and increased circumduction to ensure adequate toe clearing [40,42]. Interestingly, except in rare cases, our subjects showed no tendency toward either of the traditional kinematic compensations associated with SKG. The treadmill speeds were such that despite the limited of knee and hip ROMs they were able to achieve enough toe clearance to maintain limb motion at the given speeds that matched the treadmill belt speed. This is consistent with a recent report that ambulatory females with RTT were able to walk on a treadmill [43]. However, this report did not indicate the speeds at which their subjects walked only that they walked for six minutes at their ‘maximal’ speed. As previously mentioned, for the vast majority of our subjects as the treadmill speed exceeded 0.5 m/s our subjects exhibited signs of discomfort and the treadmill speed was immediately decreased and testing discontinued. We hypothesize that, unlike those with CP or post-stroke who walk faster than our subjects and demonstrate compensatory kinematic strategies, our subjects with RTT were unable to modify their kinematic strategies that would enable them to walk at faster speeds. Possible factors that may contribute to our participants’ slow gait speeds are discussed below.

There are several factors identified in the literature that are related to severely reduced lower limb ROMs, particularly that of the knee. Often individuals with CP and post-stroke demonstrate SKG and this is often been attributed to hyperactivity of the rectus femoris [36,44]. Another suggested cause of SKG is a lack of adequate push off at the ankle [45], leading to a reduced knee velocity at toe off and therefore reduced passive knee flexion [46,47]. Reduced hip joint velocity associated with weak hip flexors is also suggested to be a potential cause of SKG [48,49]. All of these muscle-related issues are likely to be factors in the severely reduced lower limb ROMs and contribute to slow walking speeds observed in the current study

Besides resulting in gait kinematics that significantly reduce the speed at which our subjects could walk, these kinematic patterns are energy inefficient [40,50] with oxygen consumption and cost being elevated [51]. An investigation of 12 females with RTT who walked for six minutes on a treadmill, reported that energy production was low relative to healthy subjects that could result in tiredness within a few minutes of walking [43]. As observed in Figures 3 and 4, our subjects had large symmetry indices and it is worth noting that there was a significant linear trend for knee flexion asymmetry to increase as treadmill speed increased (R^2^ = 0.90). For comparative purposes, a gait study of patients with peroneal nerve palsy displayed median knee joint angular asymmetry of 20% from perfect symmetry [37], while a group of healthy subjects displayed a 3.7% deviation from perfect symmetry [38]. In contrast, our knee joint asymmetries ranged from 26% at 0.2 m/s to 36% at 0.5 m/s. reflecting a high degree of asymmetry.

Significant kinematic asymmetries during gait are part of an overall pattern of lower limb motion that is energetically inefficient and will result in a rapid rate of fatigue development.

As has previously been reported, there was a significant positive relationship between our subject’s stride times and age [21]. Interestingly however, there were low (hip) and negligible (knee) relationships between age and ROM and age and angular velocity (Table 2). Conversely, there were significant positive relationships between both knee and hip ROM with stride time and angular joint velocities with stride time. This is consistent with previous reports in that speed is a greater indicator of associated kinematic gait variables than is age [52,53] and this relationship appears to hold true for those with RTT.

The data from the current study provides evidence that a relatively large sample of ambulatory individuals with RTT are able to walk on the treadmill and modify their kinematic pattern such that they are able to increase their walking speed within a limited range. Despite kinematic patterns that lead to SKG, poor dynamic postural control and limited concentration on the walking task, we suggest that those with RTT would benefit from a physical activity program that includes regular bouts of treadmill walking [3,5,21,43]. Heart rate, cardiac vagal tone, mean arterial blood pressure and cardiac sensitivity to baroreflex, and transcutaneous partial pressures of oxygen sampled in females with RTT respond to treadmill walking in patterns that are similar to those of healthy individuals [43]. In a recent review article focused on evaluating post-stroke physical activity programs, it was reported that three studies that used a treadmill walking intervention found significant improvements in peak oxygen uptake after the intervention [54]. These findings strongly suggest that ambulatory patients with RTT can achieve improved physical fitness resulting from a walking fitness program despite the challenges they must overcome.

Besides improved physical fitness, a second benefit of a treadmill walking program would be potential improvements in gait kinematics and postural control dynamics that could result in increases in walking speed. Increases in walking speed have been reported to improve gait kinematics. For example, 20 post-stroke subjects were exposed to a treadmill walking protocol that required them to walk as fast as possible. The results demonstrated that compared with their self-selected speed, walking as fast as possible improved the symmetry between their hemi-paretic and nonparetic limbs, as well as increases in knee and hip ROM [55]. Willerslev-Olsen, et al. [56] reported that the benefits of daily treadmill training over one month with 16 children with CP included, significant increases in speed, improved dorsiflexion during the late portion of the swing phase and increase weight acceptance on the heel during early stance. The authors proposed that treadmill gait training may promote plasticity in the corticospinal tract driven by sensory input into the CPG and results in their observed improvements in gait. Similar results following treadmill gait training were reported in patients who had incomplete spinal cord injuries [57].

An important finding is that improvement in gait kinematics can be achieved by walking at less than an individual’s maximal speed during treadmill training [55]. Although this study was completed with individuals with chronic stroke, it has direct relevance for those with RTT who often struggle to sustain their maximal achievable gait speed, even during treadmill walking. Beyond, improvement in physical fitness and gait parameters, regular walking has the potential to positively influence quality of life and wellbeing of those with RTT. Given the above information, it is reasonable to hypothesize that ambulatory females with RTT will also benefit from a treadmill gait training protocol.

In conclusion, our investigation has demonstrated that ambulatory females with RTT are able to adapt their stride times and lower limb kinematics in response to increases in treadmill belt speed, albeit within a very narrow range of gait speeds. Additionally, we have characterized several kinematic parameters associated with RTT, including very limited knee and hip ROM and significant asymmetrical motion. Despite the altered gait characteristics, we propose that a treadmill walking training program can improve the overall physical fitness as well as kinematic parameters, thereby improving the quality of life for those with RTT.

